# Plan-based generalization shapes local implicit adaptation to opposing visuomotor transformations

**DOI:** 10.1101/361634

**Authors:** Raphael Schween, Jordan Taylor, Mathias Hegele

**Affiliations:** Justus-Liebig-University, Giessen, Germany, Department of Sport Science, Neuromotor Behavior Laboratory; Princeton University, Department of Psychology, Intelligent Performance and Adaptation Laboratory

**Keywords:** motor learning, implicit, explicit, sensorimotor, dual adaptation

## Abstract

The human ability to use different tools demonstrates our capability of forming and maintaining multiple, context specific motor memories. Experimentally, this ability has been investigated in dual adaptation, where participants adjust their reaching movements to opposing visuomotor transformations. Adaptation in these paradigms occurs by distinct processes, i.e. the development of explicit aiming strategies for each transformation and/or the implicit acquisition of distinct visuomotor mappings. The presence of distinct, transformation-dependent aftereffects has been interpreted as support for the latter. Alternatively, however, distinct aftereffects could reflect adaptation of a single visuomotor map, which is locally adjusted in different regions of the workspace. Indeed, recent studies suggest that explicit aiming strategies direct *where* in the workspace implicit adaptation occurs.

Disentangling these possibilities is critical to understanding how humans acquire and maintain separate motor memories for different skills and tools. We therefore investigated generalization of explicit and implicit adaptation to different directions after participants practiced two opposing cursor rotations, which were associated with separate visual workspaces. Whereas participants learned to compensate opposing rotations by explicit strategies that were specific to the visual workspace cue, aftereffects were not sensitive to visual workspace cues. Instead, aftereffects displayed bimodal generalization patterns that appeared to reflect locally limited learning of both transformations. By varying target arrangements and instructions, we show that these generalization patterns are consistent with implicit adaptation that generalizes locally around (explicit) movement plans associated with opposing visuomotor transformations. Our findings show that strategies can shape implicit adaptation in a complex manner.

**New & Noteworthy:** Visuomotor dual adaptation experiments have identified contextual cues that enable learning of separate visuomotor mappings, but little is known about the underlying representations of learning. We report that visual workspace separation as a contextual cue enables participants to compensate opposing cursor rotations by a combination of explicit and implicit processes: Learners developed context-dependent explicit aiming strategies while an implicit visuomotor map represented dual adaptation independent from context by local adaptation around the explicit movement plan.

## Introduction

Modern tools frequently require their users to operate under altered visuomotor transformations. The fact that humans can switch between different such tools without apparently having to relearn each transformation each time has been taken as evidence for separate memories of different visuomotor transformations that can be retrieved based on context. This remarkable ability may have been fundamental to the advancement of our species (Stout and Chaminade 2007; Stout et al. 2008; McDougle et al. 2016).

Dual-adaptation paradigms have served as a useful tool to study this ability. In these paradigms, participants learn to compensate opposing visuomotor transformations, such as visuomotor cursor rotations (Cunningham 1989) or force fields (Shadmehr and Mussa-Ivaldi 1994), in close temporal succession (Bock et al. 2005; Galea and Miall 2006; Woolley et al. 2007, 2011; Hegele and Heuer 2010; Thomas and Bock 2012; Ayala et al. 2015; van Dam and Ernst 2015). Whereas alternating exposure to opposing transformations leads to substantial interference in many situations (Donchin et al. 2003; Woolley et al. 2007; Howard et al. 2013; Sheahan et al. 2016), researchers have identified a limited set of contextual cues that can enable simultaneous learning of opposing transformations (Seidler et al. 2001; Imamizu et al. 2003; Osu et al. 2004; Hegele and Heuer 2010; Howard et al. 2012, 2013; Ayala et al. 2015; Sarwary et al. 2015; Sheahan et al. 2016; Nozaki et al. 2016; Heald et al. 2018).

When it occurs, dual adaptation has been explained as contextual cues establishing separate motor memories or visuomotor mappings (Osu et al. 2004; Imamizu et al. 2007; Hirashima and Nozaki 2012; Ayala et al. 2015), or as opposite learning within a single visuomotor map being enabled by local generalization around different kinematic properties of the movement, like its trajectory (Gonzalez Castro et al. 2011), the direction of a visual target (Woolley et al. 2007, 2011), or the movement plan (Hirashima and Nozaki 2012; Sheahan et al. 2016). Whereas the privileged role attributed to movement characteristics may be justified by the omnipresence of these physical cues in natural environments, the above views can be unified by thinking of the motor memory that results from learning as a multidimensional state space that can contain arbitrary psychological and physical cue dimensions (Howard et al. 2013). Under this view, whether or not a cue enables dual adaptation depends on whether different cue characteristics allow for a regional separation in the state space of memory that is sufficient to reduce the overlap between local generalization of multiple transformations and thereby attenuate interference between them.

A level of complexity is added to this by recent views that propose at least two qualitatively distinct learning mechanisms in visuomotor adaptation (Taylor and Ivry 2012, 2014; Huberdeau et al. 2015; McDougle et al. 2016): On the one hand, there is an implicit process, which operates outside of awareness and learns from sensory-prediction errors (Mazzoni and Krakauer 2006; Synofzik et al. 2008; Morehead et al. 2017). This *implicit* learning is thought to reflect cerebellum-dependent adaptation of internal models (Taylor et al. 2010) and to dominantly contribute to aftereffects that persist in the absence of the novel transformation (Heuer and Hegele 2008). On the other hand, learners can develop conscious aiming strategies to augment reaching performance, a process referred to as *explicit* learning (Heuer and Hegele 2008; Taylor et al. 2014). This explicit learning appears to be remarkably flexible, is strongly biased by visual cues, and verbal instruction, but does not lead to aftereffects (Taylor et al. 2014; Bond and Taylor 2015, 2017).

One way to think of these distinct learning mechanisms in dual adaptation is to propose that they create two separate memory spaces that may incorporate different contextual cue dimensions, respectively. Evidence in favor of such a distinction comes from findings indicating that explicit and implicit learning differ with respect to their generalization properties (Heuer and Hegele 2011; McDougle et al. 2017) and with respect to the cue characteristics that enable dual adaptation in these two domains (Hegele and Heuer 2010; van Dam and Ernst 2015). Importantly, recent findings have also pointed towards an interaction of explicit and implicit learning mechanisms suggesting that the explicit movement plan is the center of implicit generalization (Day et al. 2016; McDougle et al. 2017; Morehead et al. 2017) and, thus, separate movement plans attenuate interference in dual adaptation (Hirashima and Nozaki 2012; Sheahan et al. 2016, 2018). Translated to our framework, this suggests that the planned movement direction (i.e. the output of the explicit memory), constitutes a relevant dimension in the state space of implicit learning. The existence of such a link would be essential to our understanding of dual adaptation and the interplay of different learning mechanisms. However, this plan-based dual adaptation is in direct conflict with previous accounts that suggested the visual target (Woolley et al. 2011) or the movement kinematics (Gonzalez Castro et al. 2011) constitute the relevant separating features.

These issues are central to our ability to use tools or different motor behaviors in different contexts. To gain more insight into the way explicit strategies and implicit learning interact in visuomotor dual adaptation, the present study seeks to test whether the direction of explicit movement plans indeed enables local learning of opposing cursor rotations by separating generalization and contrast this possibility with a previous view under which the direction of the visual target is the relevant cue (Woolley et al. 2011). We chose visual workspace as a contextual cue that has been shown to create separate explicit strategies, but not implicit visuomotor maps (Hegele and Heuer 2010) and tested generalization of explicit and implicit learning to different directions after practice. By varying target locations, rotation directions, and verbal instruction of strategies, we tested if generalization centered around the visual target (target-based generalization; Woolley et al. 2011) or the explicit strategy (plan-based generalization; Day et al. 2016; McDougle et al. 2017) limits interference within a single, implicit visuomotor map.

## Materials and Methods

A total of 94 participants signed informed consent as approved by the local ethics committee. Participants included in the analyses were right-handed and had not previously participated in a reaching adaptation experiment. Table 1 provides further information on participants and exclusions.

**Table 1:** Overview of included and excluded participants and trials for all experiments. Reasons for excluding participants were: ^a^ did not finish testing (due to bug in the experimental software or time constraints), ^b^ did not meet inclusion criteria (age, handedness, metal implants), ^c^ failure to follow instructions (revealed in post-experimental standardized questioning). Individual trials were excluded if no start could be detected or participants failed to reach target amplitude within the specified time (‘movement’) or if the angular cursor error was greater than 120° (‘angle’).

| Exp. | N analyzed | males | age (mean, min, max) | N excluded | # exclusions (movement) | # exclusions (angle) | total # of movements |
| --- | --- | --- | --- | --- | --- | --- | --- |
| 1 | 22 | 7 | 25 (20; 30) | 1<sup>a</sup> + 1<sup>b</sup> | 462 (5.5 %) | 12 (0.1 %) | 8360 |
| 2 | 20 | 4 | 21 (19; 30) | 0 | 353 (4.1 %) | 56 (0.6 %) | 8680 |
| 3 | 19 | 7 | 24 (19; 29) | 1<sup>c</sup> | 401 (4.9 %) | 152 (1.8 %) | 8246 |
| 4 | 21 | 9 | 24 (20; 29) | 1<sup>a</sup> + 1<sup>b</sup> + 7<sup>c</sup> | 320 (3.5 %) | 12 (0.1 %) | 9114 |

### Apparatus

Participants sat 1 m in front of a vertically propped 22-inch LCD screen (Samsung 2233RZ) running at 120 Hz (Figure 1A). Their index finger was strapped to a plastic sled (50 × 30 mm base, 6 mm height) that moved with low friction on a horizontal glass surface at table height. Sled position was tracked at 100 Hz by a trakSTAR sensor (Model M800, Ascension Technology, Burlington, VT, USA) mounted vertically above the fingertip. Hand vision was occluded by a black wood panel 25 cm above the surface. With their finger movement, participants controlled a cursor on the screen (cyan filled circle, 5.6 mm diameter) via a custom script written in Matlab (MATLAB, RRID:SCR_001622) using Psychophysics toolbox (Brainard 1997; RRID:SCR_002881). On *movement practice trials*, participants had to “shoot” the cursor from a visual start (red/green outline circle, 8 mm diameter) through a target (white filled circle, 4.8 mm diameter) by a fast, uncorrected movement of their right hand. Cursor feedback was provided concurrently but was frozen as soon as participants passed the target amplitude. The cursor veridically represented hand position during all familiarization and baseline practice trials (phase explanations below). During rotation practice and maintenance (inserted in-between posttests), the cursor was rotated around the start relative to hand position. The direction of cursor rotation was cued by the location of display on the screen (see below). On *movement test trials*, the cursor disappeared upon leaving the start circle. If participants took longer than 300 ms from leaving the start circle to reaching target amplitude, the trial was aborted and an error message was displayed (“Zu langsam!”, i.e. “Too slow!”). After the end of the reaching movements, arrows at the side of the screen guided participants back to the start location without providing cursor feedback.

**Figure 1:**
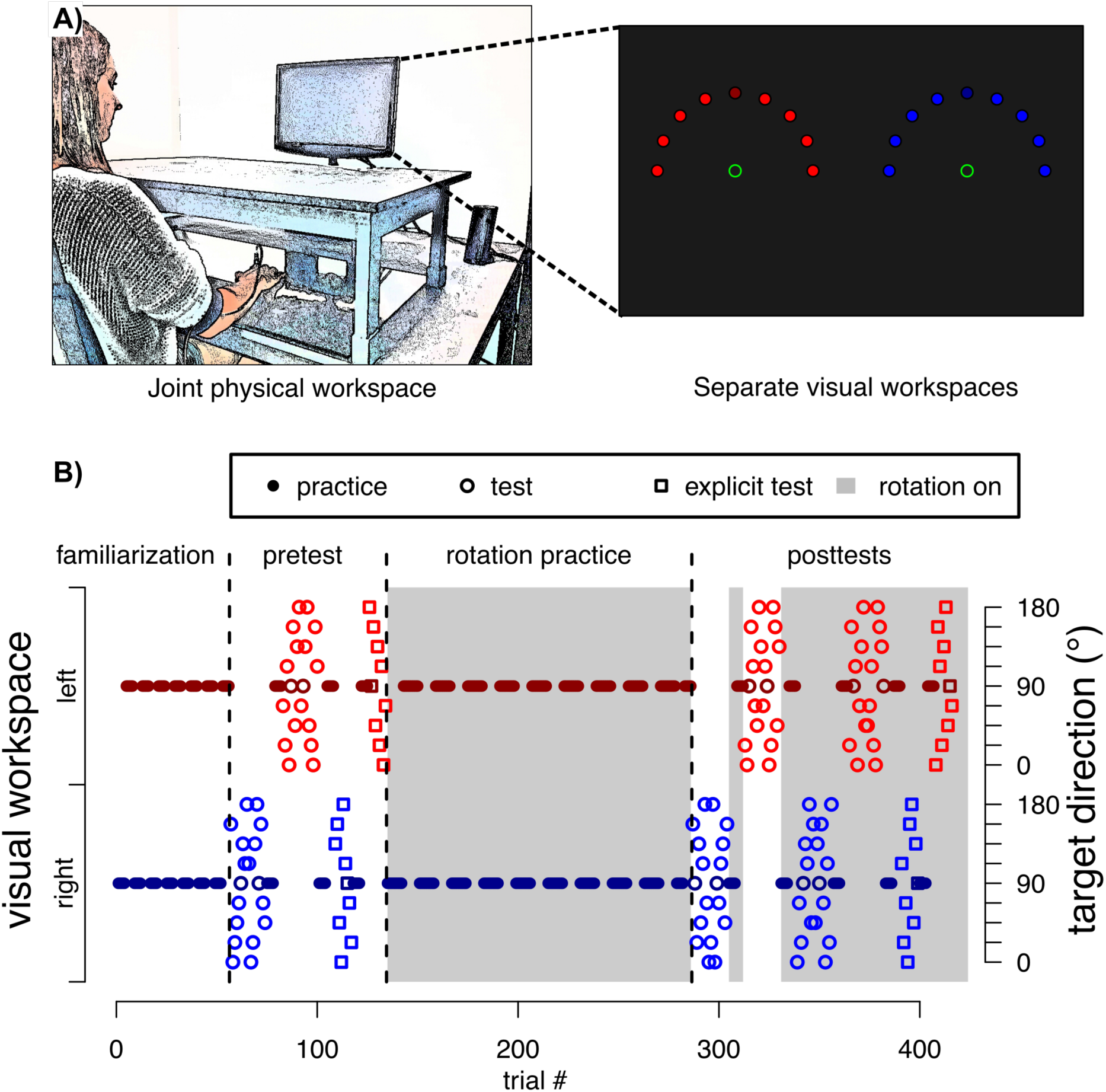
A) The general setup and visual workspace. Only one start and target location were shown on a given trial, but all generalization targets are displayed here for illustrative purposes. In addition, the actual targets were white. B) Experimental protocol for an exemplary participant of experiment 1. The start location of the hand on the table was identical for both visual workspaces. The presence/absence of the rotation was cued by the color of the start circle and instructed for both, trials with and without feedback. Alternation between visual workspaces was every four trials during familiarization and posttests practice and every eight trials during rotation practice.

### Visual workspace cue

Throughout the experiment, the start locations of the reaching movement alternated between the left and right half of the screen (x-axis shift of ¼ screen width in respective direction). In phases with cursor rotation, these visual workspaces were associated with the cursor rotation sign. We chose this contextual cue because previous research had indicated that it successfully cues separate explicit strategies but not separate, implicit visuomotor maps (Hegele and Heuer 2010). Importantly, participants’ actual movements were always conducted in a common physical workspace from a central start location on the table approximately 40 cm in front of them.

### General task protocol

All experiments consisted of familiarization, baseline pretests, rotation practice and posttests, with the general logic that posttests tested generalization of learning induced by rotation practice relative to baseline pretests. During familiarization, participants performed a total of 48 movement practice trials, with the visual workspace alternating between left and right half of the screen every four trials. This was followed by pretests intermixed with additional practice trials: in pretests, we tested generalization of movements to nine different directions spanning the hemisphere around the practiced direction by movement test trials without visual feedback. These were performed in each visual workspace in an alternating, blocked fashion (figure 1B). Each test block contained two (experiment 1) or three (experiment 2-4) sets of one reaching movement per target direction. The sequence of targets was randomized within sets. Before each test block, participants performed four more practice movements with cursor feedback in each visual workspace, respectively, to maintain reaching performance on a stable level.

At the end of pretests, we further probed participants’ explicit knowledge of the cursor rotations for one set of targets in each visual workspace. On these *explicit judgment test* trials, participants rested their hand on their thigh and provided perceptual judgments about the appropriate aiming direction to reach a specific visual target by verbally instructing the experimenter to rotate the orientation of a straight line originating on the start circle (Heuer and Hegele 2008). They were instructed that the orientation of the line should point in the aiming direction of their hand movement, which would be required to move the cursor from the start to the respective target location.

With the start of rotation practice, two oppositely signed cursor rotations were introduced and participants trained to counteract these rotations in alternating blocks of eight trials for a total of 144 trials. Before this practice phase, participants were instructed that the mapping of hand to cursor movement would be changed, that the change would be tied to the visual workspace, and that its presence would be signaled by a red (instead of the already encountered green) start circle. No further information about the nature of the change was provided.

Posttests were arranged like pretests, with few exceptions: we now repeated the movement tests twice, with the first repetition testing for generalization of *implicit aftereffects* in the absence of strategies. This was done by instructing participants before the test session, that the cursor rotation would be removed and that this would be signaled by a green start circle. The second repetition then tested for generalization of *total learning* by instructing participants that the cursor rotation was present again (as indicated by the red start circle) and they should move accordingly.

We conducted posttests for *explicit judgments* with a red start circle (transformations present) only. If participants judged the rotation to be zero at the first explicit posttests trial, they were reminded of the presence of the rotation and asked whether they wanted to reconsider their explicit judgment. This reminder was provided only once; thereafter, explicit judgments were recorded as given without further questioning.

### Experimental Protocol: Experiment 1

In experiment 1, participants practiced reaching movements to a single target direction at 90° (with 0° corresponding to movements to the right). Cursor feedback was rotated around the start location with a clockwise (CW) rotation in the left and a counterclockwise (CCW) rotation in the right visual workspace (figure 2A). Movements thus had a common visual target direction but the solutions to the two rotations were separate. Each movement generalization test block contained two sets of trials to nine equally spaced generalization targets from 0° to 180°. Pretests thus contained 86 trials, including 36 movement test trials, 18 explicit test trials and a total of 32 movement practice trials. Similarly, posttests contained 138 trials in total.

**Figure 2:**
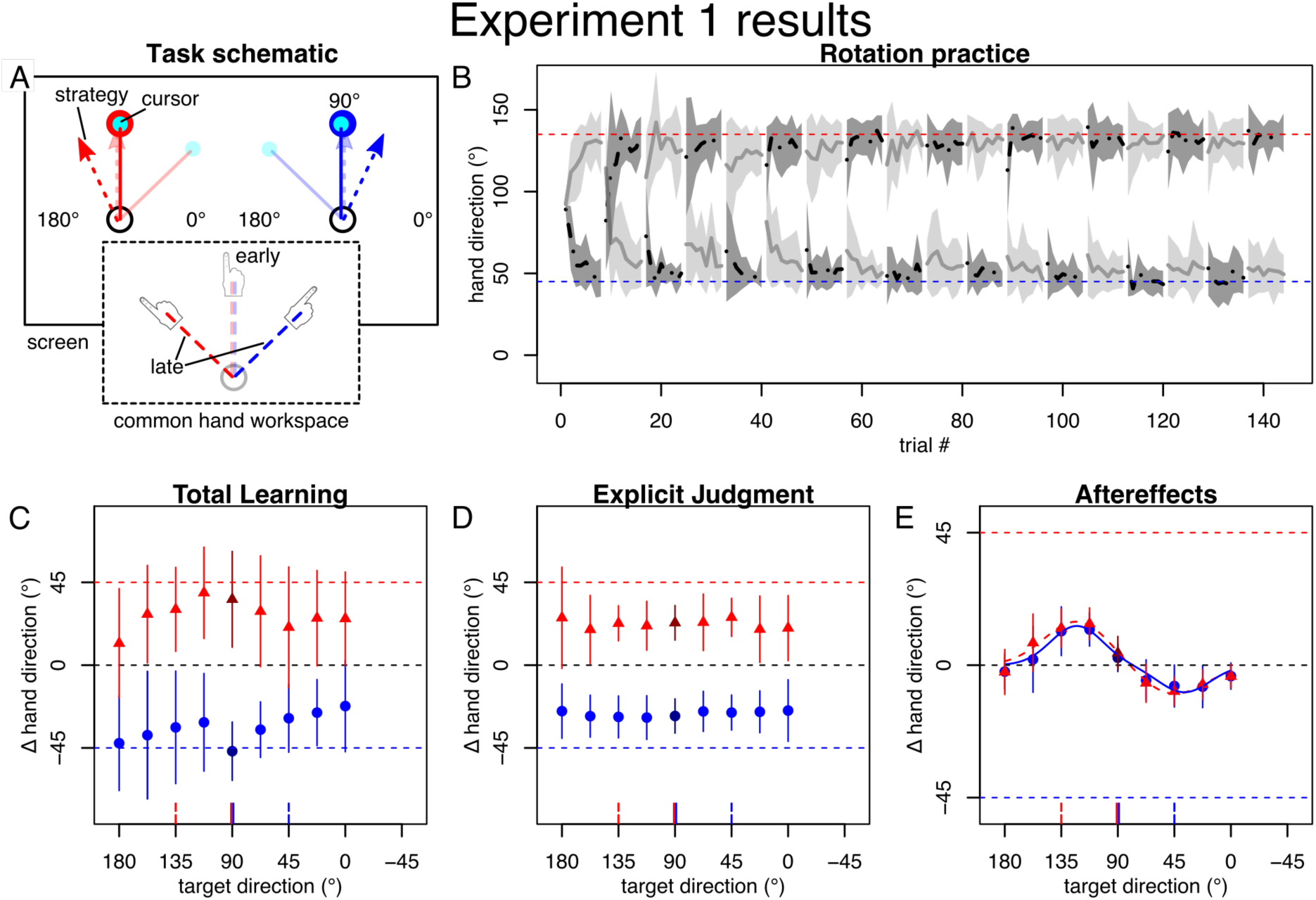
Task and results of experiment 1. All angles are in degrees with 0° falling on the x-axis and positive direction counterclockwise. Error regions and bars represent standard deviations. A: Schematic drawing of the practice targets and approximate predictions for cursor and strategy directions (top) and movement directions in the common hand workspace (bottom), for early (transparent) and late (solid) practice, for the left (red) and right (blue) visual workspace. B: Mean hand directions during practice plotted separately by the subgroups starting with either with left (grey) or right (black) visual workspace. Horizontal, dotted lines indicate ideal compensation of the cursor rotation. On average, participants compensated the rotation well within the first few blocks. C-E: Baseline-corrected average hand directions for left (red) and right (blue) visual workspace on generalization posttests. Darker color indicates the practiced target direction. Horizontal, dashed lines indicate full compensation of the cursor rotation for left (red) and right (blue) visual workspace, respectively. Vertical red and blue lines at x-axis indicate direction of target (solid) and full compensation (dashed). C: Participants’ total learning approached full compensation, was specific to the visual workspace, and generalized broadly across target directions. D: When tested separately, explicit knowledge reflected this cue-dependent, broadly generalizing learning. E: Aftereffects on the other hand appeared independent of the visual workspace cue and exhibited a generalization pattern that was well fit by a sum of two Gaussians (solid red and blue lines).

### Experiment 2

The goal of experiment 2 was to ensure that our findings from experiment 1 were not solely attributable to biomechanical or visual biases independent of learning (Ghilardi et al. 1995; Morehead and Ivry 2015). To test this possibility, the practice and generalization targets were moved by 45° CW (i.e. the practice target was at 45° and generalization targets spanned -45° to 135°; Figure 3A). We predicted that if the generalization pattern was solely due to potential biases, then it would be unchanged. However, if the apparent generalization function was the result of learning, then it should be shifted by -45° (i.e. 45° CW) on the generalization direction axis. Apart from these changes, experiment 2 was like experiment 1, except that we increased the number of consecutive movement test sets for each visual workspace and test condition from two to three, thus increasing the number of pretest trials to 122 and posttest trials to 174.

**Figure 3:**
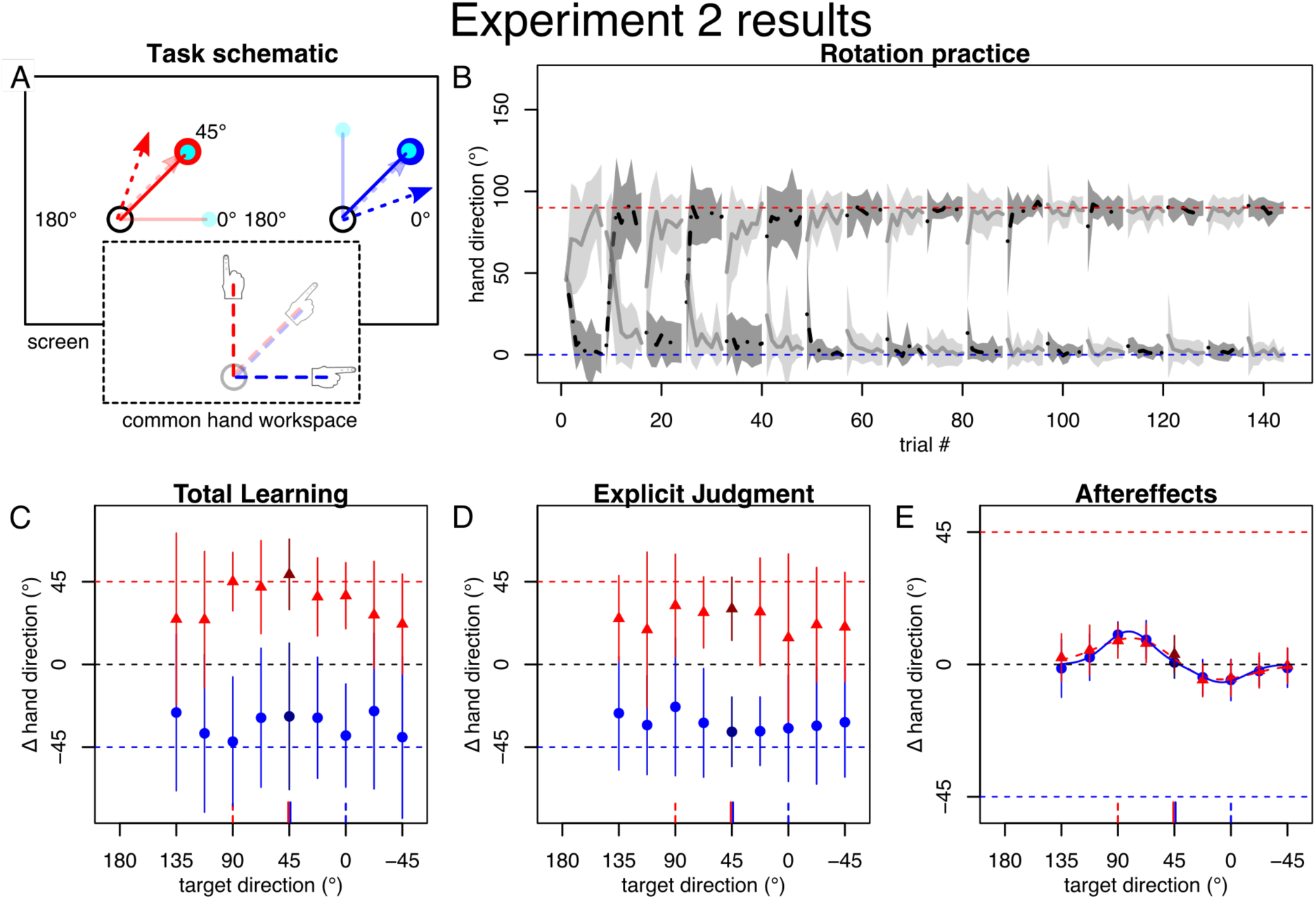
Task and results of experiment 2. A: Practice direction was rotated by -45° relative to experiment 1. B-D: Participants quickly learned to compensate the rotation and displayed appropriate total learning and explicit knowledge, as in experiment 1. E: Generalization of aftereffects appeared shifted with the practice location, indicating that bimodal pattern is dominantly an effect of learning, not biases.

### Experiment 3

To further contrast plan-based and target-based generalization, we designed a paradigm with separate visual target locations and cursor rotations, which were arranged in such a way that the resulting compensation strategy for each target should approximately point at the respective other target when projected to the common physical workspace. This way, plan- and target-based generalization predict opposite generalization patterns. We therefore offset targets by 22.5° outward from the center (i.e. to 112.5° in the left and 67.5° in the right workspace). For rotations, we chose 60° CCW for the left and CW for the right workspace (figure 4A). Since implicit learning asymptotes at about 10-15° regardless of rotation magnitude (Bond and Taylor 2015; Morehead et al. 2017), we assumed that strategy magnitude should asymptote around the 45° inter-target distance, as intended. Otherwise, experiment 3 was identical to experiment 2.

**Figure 4:**
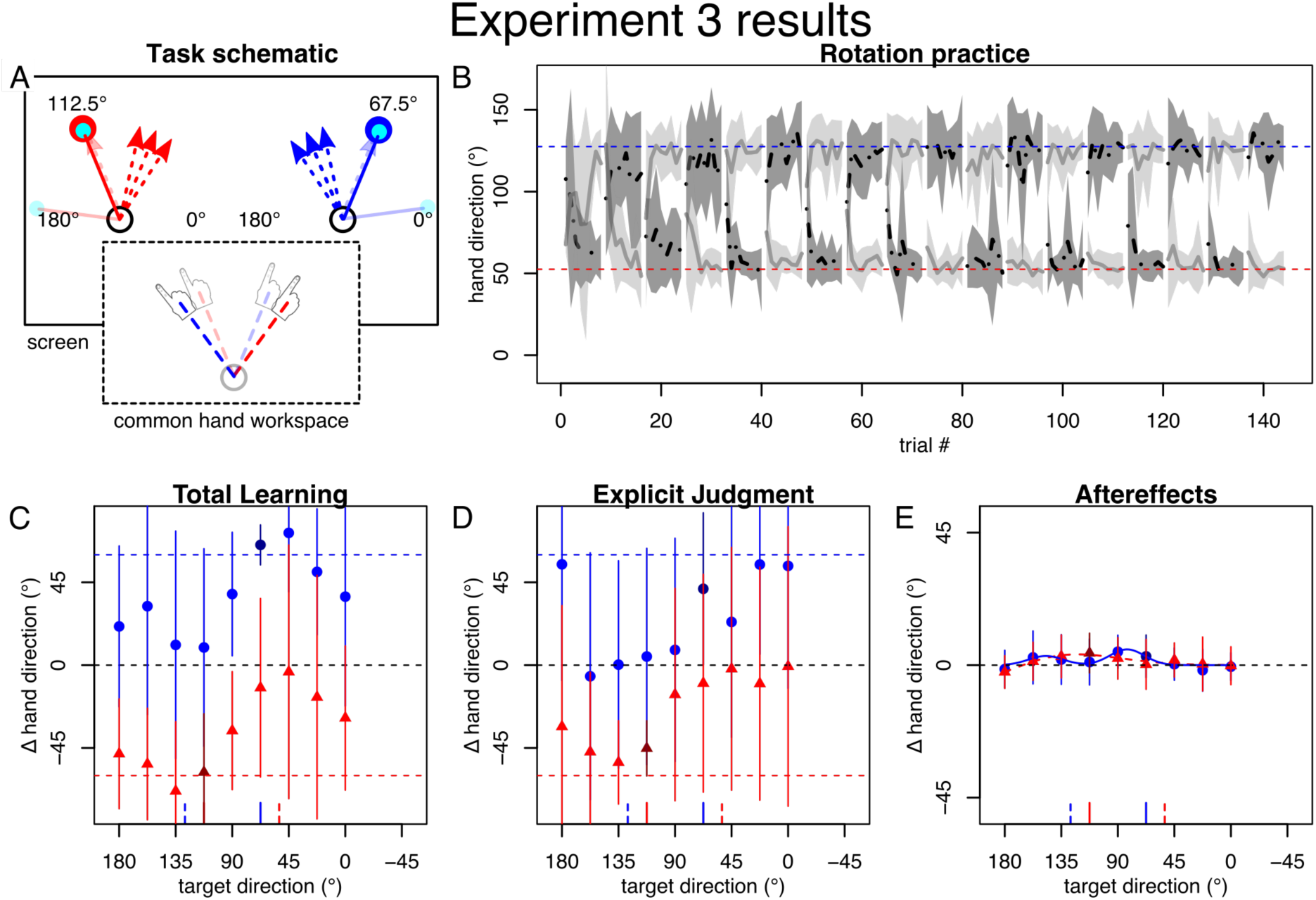
Task and results of experiment 3. A: Visual target directions were separated by 45° and the cursor rotated outward so that strategies should cross the midline and point approximately at the other target. B: Practice performance was more variable compared to experiment 1 (A). C-D: Total learning and explicit judgments flipped signs in line with the reversed cursor rotations cued by the separate visual workspaces. They also appeared more variable, but resembled experiment 1 when removing 5 outliers (not shown). E: Aftereffects appeared dominated by interference, which did not change without the 5 outliers (not shown). Note that the lines indicating full compensation are outside the y-axis-limits.

### Experiment 4

This experiment intended to facilitate correct aiming strategies by verbally instructing participants about how to compensate the cursor rotation before practice of the rotations. This was done by informing participants that they would have to aim roughly toward 1 o’clock in the left workspace and 11 o’clock in the right workspace to hit the respective practice targets. They were also encouraged to fine-adjust those strategies. We hypothesized that this instruction should strengthen the contrast we originally hypothesized in experiment 3 (figure 4A). To ensure that participants applied non-overlapping strategies throughout practice, we asked participants after the experiment where they aimed during early, middle, and late practice in the left and right workspace, respectively. However, based on these post-experiment reports, we excluded 7 participants who reported not using the clock analogy or aiming less than half an hour in the correct direction away from 12 o’clock for any of those time points (table 1).

### Data analysis

Data were analyzed in Matlab (MATLAB, RRID:SCR_001622), R (R Project for Statistical Computing, RRID:SCR_001905), and jasp (JASP, RRID:SCR_015823). Position data were low-pass filtered using Matlab’s “filtfilt” command set to a 4^th^ order Butterworth filter with 10 Hz cutoff frequency. We separately calculated x- and y-velocity using a two-point central difference method and tangential velocity. Movement start was determined as the first frame where participants had left the start circle and tangential velocity exceeded 30 mm^∗^s^-1^ for at least three consecutive frames. For each trial, we extracted the angular endpoint direction as the angle between the vector from start to target and the vector between the start and the position where the hand passed the target amplitude. We excluded trials where no movement start could be detected or where participants failed to reach the target amplitude (see table 1).

For pre- and posttests, we calculated medians separately for each visual workspace (combining the groups with different initial workspaces), target direction and type of posttest. Our main variables of interest were generalization patterns of aftereffects and we limit our analysis on explicit judgments and total learning to descriptive reporting.

In our analysis of implicit learning, we used two candidate functions to represent our hypotheses for the shape of generalization. The first candidate was a single Gaussian:

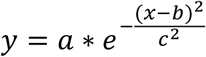

where *y* is the aftereffect at test direction *x* and the three free parameters are the gain *a*, the mean *b* and the standard deviation *c.*

The second candidate was the sum of two Gaussians, henceforth referred to as “bimodal Gaussian”:

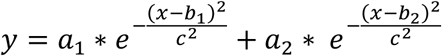

for which we assumed separate amplitudes *a*_1_; *a*_2_ and means *b*_1_; *b*_2_ but the same standard deviation *c* for the two modes.

We reasoned that successful learning of separate visuomotor maps by visual workspace cues should result in separate generalization curves for the cue conditions where each should resemble a single Gaussian in a direction appropriate for counteracting the cued rotation. Note, we did not expect such an outcome based on previous results (Hegele and Heuer 2010). If the cues did not establish separate visuomotor maps on the other hand, we predicted either of two patterns for the resulting, common generalization curves. Depending on whether the centers of local generalization were overlapping or separate, the pattern should be dominated by interference, which could be fit by either of the functions but with small amplitude parameters, or by a bimodal generalization pattern with opposing peaks whose centers and amplitudes would be in line with compensating the opposing cursor rotations. Which scenario would be the case depends on whether generalization is target-based (Woolley et al. 2011) or plan-based (Day et al. 2016; McDougle et al. 2017), as well as on the specific arrangements of plan and strategic solutions in the different experiments. We could further have included a single Gaussian with an offset parameter in our model comparison but decided against this option as our focus was to distinguish between types of dual adaptation rather than to infer its exact shape.

To test our hypotheses, we fit the two candidate models to the nine group mean data points corresponding to the nine generalization directions of each visual workspace’s aftereffect, using Matlab’s “fmincon” to maximize the joint likelihood of the residuals. For this, we assumed independent, Gaussian likelihood functions centered on the predicted curve, whose variance we estimated by the mean of squared residuals. As this fitting procedure tended to run into local minima, we repeated each fit 100 times from different starting values selected uniformly from our constraint intervals (constraints were -180° to 180° on *a*, 0° to 180° on b-parameters, or 135° to -45° for experiment 2, and 0° to 180° on *c)*, and used only the fit with the highest joint likelihood.

To select the best model, we calculated Bayesian Information Criterion (BIC) as:

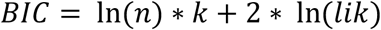

where *n* is the number of data points, *k* is the number of free parameters of the model and *lik* is the joint likelihood of the data under the best fit parameters.

To compare model parameters, we created 10000 bootstrap samples by selecting *N* out of our *N* single participant datasets randomly with replacement and taking the mean across participants for each selection. We then fit our candidate models to each of these means by the same method described above, except that we avoided restarting from different values and used the best fit values from the original dataset as starting values instead. Because the bimodal Gaussian has two identical equivalents for each solution, we sorted the resulting parameters so that *b*_1_ was always larger than *b*_2_. This procedure gave us a distribution for each parameter from which we calculated two-sided 95% confidence intervals by taking the 2.5^th^ and 97.5^th^ percentile value. We considered parameters significantly different from a hypothesized true mean if the latter was outside their 95% confidence interval. Similarly, we considered differences between two parameters significant if the 95% confidence interval of their differences within the bootstrap repeats did not include zero. Additionally, we used t-tests to compare aftereffect magnitudes (of the raw, non-bootstrapped data) for specific generalization directions against 0 or between groups. To protect ourselves from interpreting differences between experiments purely based on the absence of an effect in one and presence in the other (Nieuwenhuis et al. 2011), we further performed an ANOVA on aftereffect posttests with the between-participant factor of experiment and the within-participant factors of visual workspace and target direction, where we added 45° to the target directions of experiment 2.

## Results

We report angles in degrees with 0° corresponding to the positive x-axis and higher angles being counterclockwise with respect to lower ones. Values are reported as mean with standard deviation (SD) or with 95% confidence interval.

### Experiment 1

Participants appeared overall able to compensate for the two opposing rotations after few blocks of practice (fig 2B). Posttests for *total learning* also indicate that participants learned to compensate for the opposing rotations almost completely and specific to the visual workspace cue (figure 2C) with mean total learning towards the practiced target location falling somewhat short of full 45° compensation by compensating 36° (SD 26) in the left and even slightly exceeding the full -45° compensation by compensating -47° (SD 16) in the right workspace. Total learning tested at generalization target locations tended to generalize broadly, appearing relatively flat across directions. Explicit learning was also specific to the workspace with mean explicit judgments at the practice target amounting to 23° (SD 9) relative to 45° full compensation for the left and -28° (SD 9) relative to -45° full compensation for the right workspace and tended to display broad generalization (figure 2D).

For aftereffects (figure 2E), target-based generalization predicted separate, single Gaussians with peak directions reflecting learning of the cued workspaces if the visual workspace cue enabled the formation of separate visuomotor maps; or if the cues failed to enable global dual adaptation, then no clear peaks would be observed. Plan-based generalization, on the hand, predicts a generalization function with two opposite peaks corresponding to participants’ strategic aims.

Model comparison preferred the bimodal Gaussian for characterizing the data, as indicated by differences in BIC greater than ten (ΔBIC: 16 for left, 14 for right workspace cue) relative to the single Gaussian, which is considered “strong” evidence against the model that has the higher BIC (Kass and Raftery 1995). The amplitude parameters had opposing signs and their confidence intervals did not include zero (left: a_1_: 14.9° [12.8°; 17.4°], a_2_: -9.9° [-11.9°; - 8.5°]; right: a_1_: 13.3° [11.2°; 46.8°], a_2_: -9.2° [-22.0°; -6.6°]). The corresponding means were located roughly where we would have expected aiming strategies to lie (left: b_1_: 122.5° [117.1°; 127.9°], b_2_: 45.7° [38.8°; 55.6°]; right: b_1_: 122.6° [117.0°; 126.3°], b_2_: 37.1° [25.7°; 53.4°]; c-parameter left: 36.9° [31.1°; 44.3°]; right: 29.1° [15.9°; 39.3°]), although individual variability and the lack of separate time series for implicit and explicit learning confined us to qualitative analyses in this respect. Mean aftereffects at the practiced target were small, albeit significant (left: 4° (SD 6), *P* = .002, right: 3° (SD 5), *P* = .02), indicating that interference dominated here. This finding argues strongly against target-based generalization.

Importantly, the curves for the left and right workspace were almost indistinguishable (Fig. 2) and the confidence intervals for differences between left and right workspace parameters all included 0° (Δa_1_: [-33.2°; 4.6°]; Δb_1_: [-6.6°; 7.7°]; Δa_2_: [-3.5°; 10.8°]; Δb_2_: [-6.1°; 22.8°]; Δc: [-4.3°; 26.1°]). It therefore appears that visual workspaces did not cue separate implicit visuomotor maps.

Overall, the observed generalization curves are well in line with dual adaptation expressed locally around the movement plan or trajectory, but do not support local generalization around the visual target or separate visuomotor maps established based on visual workspace cues.

### Experiment 2

For experiment 2, we predicted a two-peaked generalization pattern of aftereffects, similar to the one we observed in experiment 1, but shifted by 45°. That is, if the pattern in experiment 1 were just biases that did not reflect learning, we would predict it to be exactly the same as in experiment 1, whereas if it were a result of learning, we would predict it to be shifted by -45° on the x-axis, reflecting the -45° shift of the practice targets.

Practice and posttests again indicated that participants learned to compensate for the cursor rotation specific to the visual workspace (figures 3B-C). Total learning tested at the practice location was 49° (SD 19) in the left workspace and thus somewhat exceeded full compensation of the -45° cursor rotation. In the right visual workspace, it fell somewhat short of fully compensating the +45° cursor rotation, with average compensation at the practiced target amounting to -28° (SD 40). Explicit judgments were on average 30° (SD 17) relative to 45° full compensation in the left and -37° (SD 19) relative to -45° full compensation in the right workspace (figure 3D).

Model comparison favored the bimodal Gaussian to characterize the generalization pattern of aftereffects (ΔBIC: 17 for left, 19 for right workspace cue). Importantly, the generalization curves indeed appeared shifted on the x-axis with complete interference occurring close to the practice direction (figure 3E). For the right workspace, this was reflected in the bootstrapped distributions of differences between experiment 1 and experiment 2 mean parameters including -45° (Δb_1_: [-66.6°; -34.3°], Δb_2_: [-48.9°; 5.7°]) and the difference between amplitudes including zero (Δa_1_: [-12.4°; 61.4°]; Δa_2_: [-45.4°; 9.9°]). For experiment 2 the right workspace parameters and confidence intervals were: a_1_: 11.2° [9.4°; 75.6°], b_1_: 81.3° [55.6°; 85.4°]; a_2_: -6.1° [-55.1°; -3.6°]; b_2_: 6.9° [-3.9°; 44.7°]; c: 25.8° [7.5°; 49.9°]. For the left visual workspace, the best fit was achieved by a solution containing two relatively close peaks with large amplitudes and standard deviations: a_1_: 162.9° [84.7°; 163.8°], b_1_: 49.9° [42.9°; 64.3°]; a_2_: -159.8° [-161.0°; -83.2°]; b_2_: 47.4° [36.0°; 57.5°]; c: 49.9° [37.5°; 63.5°]. This fit produces the visual pattern mainly by interference. While this solution was thus not easily comparable to the two largely separate peaks of experiment 1 and the right workspace, we note that both amplitudes were still significant in opposite directions and that the switch between peaks still appeared to be around the practiced target, with aftereffects at the practiced target amounting to 3° (SD 6, P_19_ = .026) in the left and 1° (SD 5, P_19_ = .59) in the right visual workspace.

Overall, we conclude from experiment 2 that, whereas some additional biases may contribute to the results observed in experiment 1, the shape of the generalization curve first and foremost reflects learning.

### Experiment 3

While experiments 1 and 2 already favored plan-based over target-based generalization, we designed experiment 3 to maximize the contrast between the two hypotheses (figure 4A). For this purpose, we had participants practice a 60° cursor rotation to a target at 112.5° in the left and a -60° cursor rotation to a target at 67.5° in the right workspace. For this scenario, target-based generalization predicted a generalization function for aftereffects with a positive peak at 67.5°and a negative peak at 112.5°. Plan-based generalization on the other hand should create the opposite result, i.e. a negative peak close to the 67.5° target direction, reflecting compensation of the positive rotation experienced with the 112.5° target and an assumed positive compensation strategy, and a corresponding positive peak close to the 112.5° target direction.

To our surprise, aftereffects no longer displayed a clear generalization pattern as in the previous experiments, but a pattern that appeared dominated by complete interference across all directions between the opposing rotations (figure 4E). Accordingly, while mean aftereffects were still best described by bimodal Gaussians (ΔBIC left: 5; right: 4), their peaks’ locations did not match either of the hypotheses and associated amplitudes were either positive and small (right workspace: 3.1° [-29.1°; 38.4°], b_1_: 147.3° [111.4°; 180.0°]; a_2_: 5.3° [3.3°; 175.0°]; b_2_: 82.6° [71.7°; 178.7°]; c: 19.5° [26.9°; 144.4°]) or excessively large (left workspace: a_1_:-125.7° [-180.0°; 124.9°], b_1_: 179.5° [95.6°; 180.0°]; a_2_: -123.7° [-121.4°; 180.0°]; b_2_: 176.3° [93.7; 178.7°]; c: 69.6° [26.9°; 144.4°]). Note that these fits approached our bounds on a and b parameters and the amplitude parameters only differed significantly from zero for right a_2_). Overall, the fit appeared unstable as shown by the histogram of bootstrapped parameter estimates (figure 6A). The visual impression of aftereffects did not change even if we removed five participants (by visual selection) who appeared responsible for the less clearly separated patterns of explicit and total learning (figure 4C-D; data with participants removed are not shown).

How could this absence of a clear generalization pattern be explained? We hypothesized that the development of aiming strategies might not have been quick enough to allow local generalization to occur in sufficiently distinct directions, thereby creating a generalization pattern that was primarily governed by interference. An indication that this might be the case can be seen in figure 4B, where mean hand directions during practice initially fall short of the ideal hand directions, which indicates poorly developed strategies at this time. If participants made such strategy errors, then it would cause counteractive learning under the plan-based generalization hypothesis and, therefore, could explain interference in posttests. To test if this was the reason for interference in experiment 3 aftereffects, we conducted experiment 4 where we provided participants with ideal aiming strategies at the onset of rotation practice. We hypothesized that more appropriate strategy application should alleviate interference and restore the predicted, plan-based generalization pattern if our reasoning was correct.

### Experiment 4

Predictions for experiment 4 were the same as they had been for experiment 3, but here we predict more local, direction-specific implicit adaptation because participants should have a more consistent strategy. Indeed, we observed more consistent performance during initial practice and the restitution of flat overall learning and explicit judgment patterns, indicating that participants were able to implement the provided strategy (fig 5B-D). Consistent with our prediction, the resulting generalization pattern of implicit learning once again had opposite peaks (ΔBIC left: 9; right: 9; figure 5E). Parameter histograms display more confined peaks compared to experiment 3 (figure 6), suggesting that the bimodal Gaussian was more appropriate, here. Importantly, the signs of the amplitude parameters were in line with the predictions under plan-based generalization and the confidence intervals on associated amplitude parameters did not include zero (left: a_1_: 6.6° [2.8°; 59°], b_1_: 112.0° [84.1°; 122.7°], a_2_: -5.5° [-58.2°; -2.9°], b_2_: 29.7° [21.8°; 58.1°], c: 25.0° [7.0°; 47.2°]; right: a_1_: 8.7° [5.1°; 13.0°], b_1_: 118.4° [111.6°; 124.8°], a_2_: -5.7° [-8.3°; -3.5°], b_2_: 29.1° [20.5°; 39.9°], c: 29.3° [22.3°, 37.6°]). This result is in direct contrast to target-based generalization. The confidence intervals for differences between left and right parameters once again included zero (Δai: [-5.2; 49.2]; Δb_1_: [-31.0; 6.5]; Δa_2_: [-52.7; 3.3]; Δb_2_: [-9.8; 28.9]; Δc: [-24.8; 18.1]), suggesting no significant influence of the contextual visual workspace cue.

**Figure 5:**
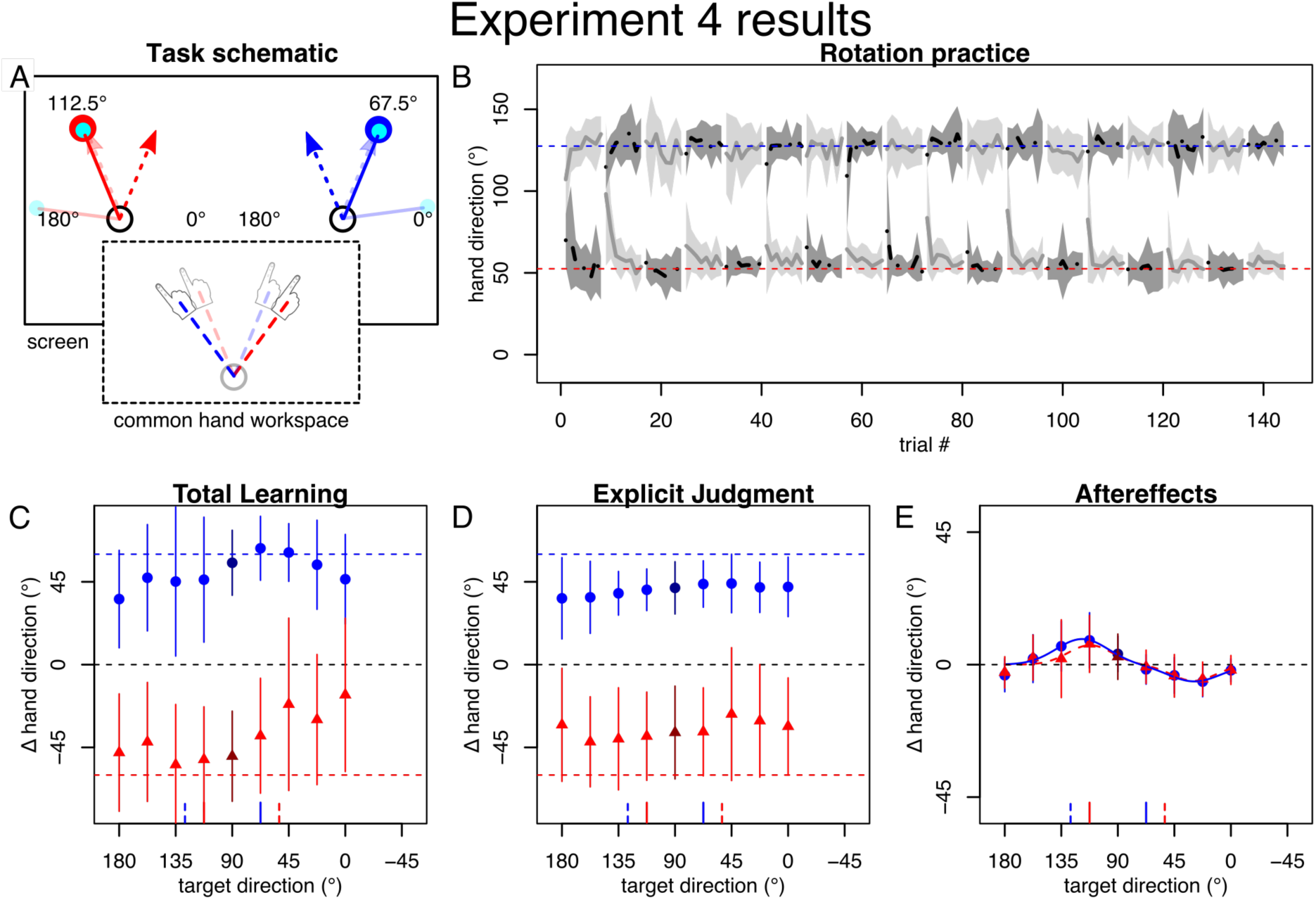
Task and results of experiment 4. A: Same scenario as experiment 3, but we provided strategies in advance. B: Practice performance appeared less variable. C-D: All participants displayed good total learning and explicit knowledge. E: The bimodal pattern of aftereffects was restored.

**Figure 6:**
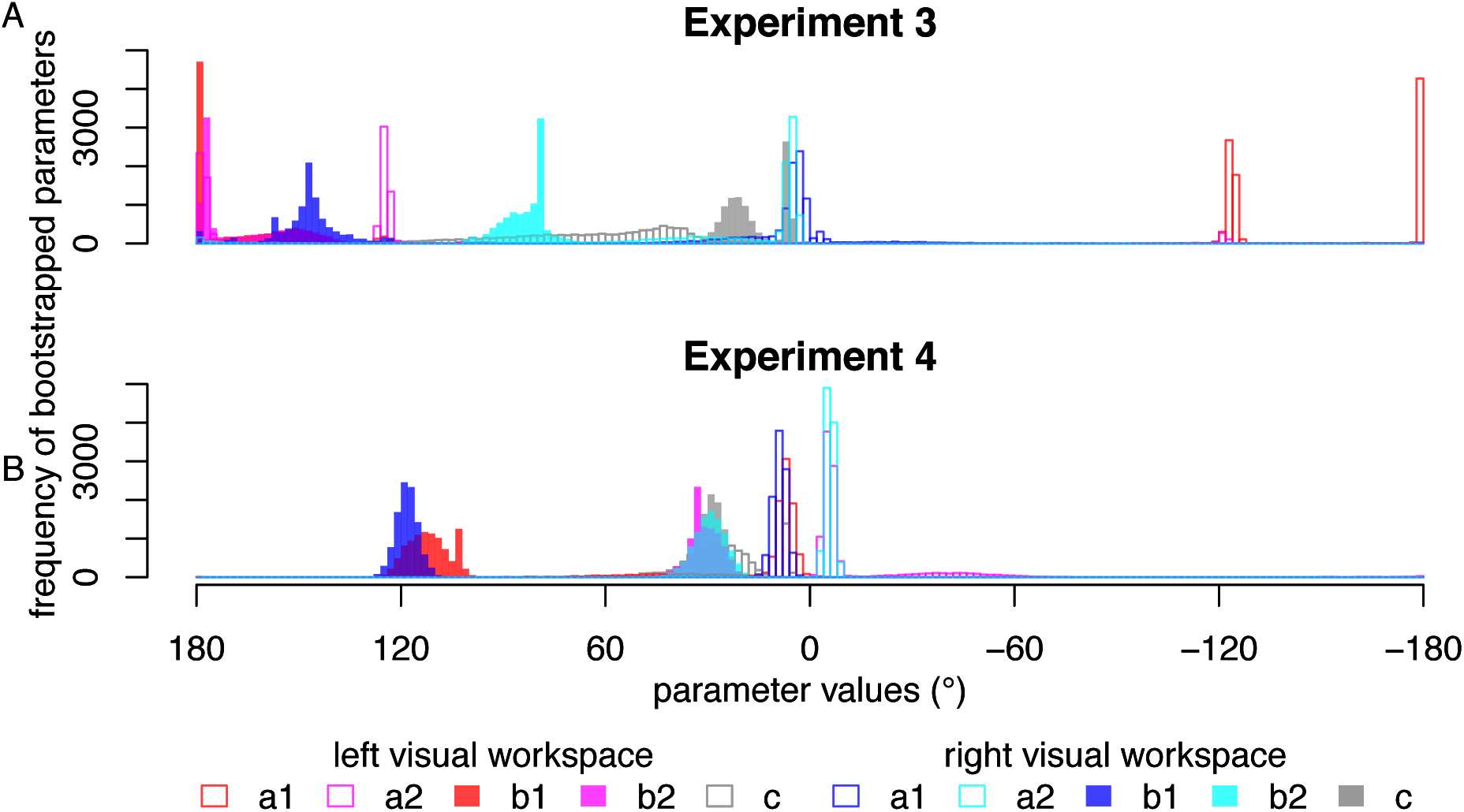
Histograms of the bootstrapped parameters for A: experiment 3 and B: experiment 4. The distributed and split peaks in A visualize the unstable fit of the bimodal Gaussian to the data of experiment 3. The more confined peaks in B show that the fit is more stable. The centers of generalization (b_1_, b_2_) prefer generalization centering on the movement plan rather than the visual target.

In comparison with experiment 3, providing an aiming strategy at the onset of practice alleviated interference in implicit learning, which we interpret to reflect better expression of local generalization due to more appropriate application of spatially separate explicit strategies.

### Across experiment comparison

To ensure that the differences we inferred from generalization patterns across experiments were statistically justified, we performed an ANOVA on aftereffect posttests with the factors experiment, workspace cue, and target direction. Greenhouse-Geisser-corrected p-values of the ANOVA across experiments indicated a significant main effect of target (*F*_3.6,207.4_ = 40.2, *P* < .001), but no other significant main effects (experiment: *F*_3,57_ = .19, *P* = .91; workspace: *F*_1,57_ = 1.5, *P* = .23). There wasn’t a significant two-fold interaction involving workspace (workspace^∗^experiment: F_3,57_ = 1.1, *P* = .38; workspace^∗^target: *F*_6.3,360.9_ = .93, *P* = .48), but a significant interaction between experiment and target direction (*F*_10.9,207.4_ = 4.5, *P* < .001). The three-way interaction approached significance (*F*_19.0,360.9_ = 1.5, *P* =.075). While being admittedly posthoc, these numbers overall support our interpretation of differences in generalization to different targets across experiments and further lend some support to the absence of a relevant influence of visual workspace cue on aftereffects.

## Discussion

The study of visuomotor dual adaptation has frequently been motivated by an interest in understanding how the brain associates contextual cues with separate memories to represent and switch between different visuomotor environments, like controlling tools or objects (Imamizu et al. 2003; Ayala et al. 2015; Heald et al. 2018). However, the underlying components contributing to this type of learning have received little attention. This has important consequences since, for example, explicitly learned behavior is subject to constraints in its expression that differ from those of implicit learning (Maxwell et al. 2001; Masters et al. 2008). Furthermore, abstract context cues may be irrelevant for separate memories when spatial constraints of practice enable opposing transformations to be accounted for within a single visuomotor map (Woolley et al. 2007, 2011; Gonzalez Castro et al. 2011; Hirashima and Nozaki 2012), a possibility that depends on the generalization properties of learning (Day et al. 2016; McDougle et al. 2017). To broaden our understanding of the underlying constituents of visuomotor dual adaptation, we investigated generalization and interference of learning when separate visual workspaces cued alternating, opposing visuomotor cursor rotations. By varying rotation size, arrangement of visual targets and instructions, we show that implicit dual adaptation is expressed as a local generalization pattern in this case. Visual workspaces cued separate aiming strategies but did not establish separate implicit visuomotor maps, thus corroborating previous suggestions based on cerebellar imaging, that separate memories for different contexts rely on cognitive components (Imamizu et al. 2003). The pattern of implicit dual adaptation we observed can be explained by generalization occurring locally around the (explicit) movement plan. Specifically, we observed peak learning at the approximate locations and in the directions predicted if we assume that learning generalizes locally around the aiming strategy, in line with recent findings (Day et al. 2016; McDougle et al. 2017). Furthermore, interference occurred in a scenario where it could be explained by generalization centering on the movement plan, but not the visual target (experiment 3).

Within the framework entertained in the introduction, the observed results strongly suggest that the planned movement direction, but not the visual workspace is a relevant dimension in the implicit memory space. Conversely, the flat pattern of explicit generalization would indicate that visual workspace, but not direction (whether plan, movement or target) was a relevant dimension in the explicit memory space. Given the high flexibility of human cognition and explicit learning, we would not expect the latter to be a general characteristic of explicit learning, though. It seems more plausible that explicit learning can account for contextual cues in arbitrary dimensions, given that learners become aware of the relevant contingencies between cues and transformations. For implicit memory on the other hand, a memory space of relatively fixed, low dimensionality would fit well with its overall simplicity, which makes it less flexible (Mazzoni and Krakauer 2006; Bond and Taylor 2015), but robust to constraints on cognitive processing (Fernandez-Ruiz et al. 2011; Haith et al. 2015). Which other cues, besides the movement plan, belong to the implicit memory space and whether more extensive practice can introduce new context dimensions to it, e.g. by associative learning as suggested previously (Howard et al. 2013), are interesting topics for future research. A practical implication of this view would be that “A-B-A” paradigms where participants learn first one (A), then another (B) and then the first transformation (A) again may be more suited to infer preexisting context dimensions, as they minimize contrastive exposure to new contingencies that could be learned associatively. Specific investigation of the latter, on the other hand might benefit from exploiting known characteristics of associative learning.

### Plan or target-based?

Our findings contradict conclusions from an earlier study, which inferred the visual target to be the relevant center of local, implicit generalization to different directions (Woolley et al. 2011). We explain this contradiction by the fact that this earlier study only compared the two alternative hypotheses that learning centers on the visual target or the executed movement but did not consider the possibility that it centers on the movement plan. When reinterpreted in the light of this new hypothesis, all results in that study can potentially be explained by plan-based generalization with separate visual targets cuing separate aiming strategies. Specifically, participants in that study learned to compensate opposing cursor rotations when visual targets were separate but ideal physical solutions overlapped. Alternatively to local generalization centering on the visual target, this can be explained by different aiming strategies becoming associated with the separate targets, each of which is less than the optimal, full rotation (Bond and Taylor 2015) and therefore does not overlap with the strategy for the opposing cursor rotation (in contrast to the physical solutions). Similarly, interference scaling inversely with the separation of visual targets (Woolley et al. 2011) may also be explained by the degree of overlap between aiming strategies.

It is worth nothing that the lack of dual adaptation in an earlier experiment by Woolley and colleagues (2007) may be attributable to the saliency of the visual cues. In this study, they found that practice to the same visual target did not enable dual learning when opposing rotations were cued by screen background colors, but the task relevancy of these cues may not have been noticed by the participants. If the participants did not associate an aiming strategy with the cues, then it would have not allowed plan-based and directionally-dependent implicit adaptation to develop; thus, leading to no dual adaptation.

### Plan or movement-based?

Alternatively to plan-based generalization, our findings could be explained by learning generalizing around the movement path, as has been found for force field adaptation (Gonzalez Castro et al. 2011). These two options are difficult to tease apart with our current methodology because the strategic movement plan deviates from the visual target in the same direction as the movement trajectory, resulting in qualitatively similar predictions for the two hypotheses. However, a number of recent findings prefer plan-based over movement-based generalization based on more specific analyses. Thus, plan-based generalization is supported by recent quantitative analyses in cursor rotations (Day et al. 2016; McDougle et al. 2017; Morehead et al. 2017). Furthermore, a recent study showed that, when participants plan to move two cursors to two separate targets by a hand movement towards the center between them, aftereffects occur locally around both targets, but interference dominates when they plan to move to the central target, instead (Parvin et al. 2018). In force fields, three recent studies showed that interference is reduced when similar trajectories are associated with different movement plans while practicing opposing force fields (Hirashima and Nozaki 2012; Sheahan et al. 2016, 2018). Similarly, opposing force fields were learned with the same trajectory when participants intended to control different points on a virtual object (Heald et al. 2018). Finally, irrespective of our reinterpretation above, Woolley and colleagues’ (2011) results show that local dual adaptation with similar physical movements is possible, providing further evidence against movement-based generalization.

Rather than a fixed center of generalization, one might expect that the brain adaptively exploits the task structure by linking memory separation to those cues that are sufficiently distinct. Our data for implicit learning do not support this possibility, as otherwise, we would have expected aftereffects in experiment 3 to be shaped by learning around the separate visual targets rather than interference. However, it is still possible that such a shift in cue relevance may occur under different circumstances (e.g. longer practice). More generally, we may ask if local learning of multiple transformations evolves according to an underlying model that is specifically adapted to the practice scenario or if it is merely the sum of single transformation learning. Previous studies have considered a model-based approach and varied practiced target directions to distinguish between these possibilities (Bedford 1989; Pearson et al. 2010; Woolley et al. 2011). We note that this approach becomes more difficult under plan-based generalization, since the centers of single adaptation can no longer be taken to be fixed but depend on flexible cognitive strategies, thus complicating quantitative inference. As noted earlier, we would expect explicit strategies to be in principle highly adaptable to even complex task structures under the right circumstances, although some default preferences may exist (Bedford 1989; Redding and Wallace 2006; van Dam and Ernst 2015), whereas implicit learning is likely more stereotypical.

### Relation to force field learning

With respect to the relevance of our findings to visuomotor transformation learning in general, we need to consider the possibility that dual adaptation in force fields may differ from that in cursor rotations. Presumably due to the less transparent nature of the transformation, aiming strategies are harder to conceptualize in force field learning and may play less of a role (McDougle et al. 2015). Furthermore, it is possible that the state space representing implicit internal models for force compensation incorporates more and different dimensions than a visuomotor map representing cursor rotations. For example, movement velocity is theoretically relevant for compensating velocity-dependent force fields, but not for velocity-independent cursor rotations.

These differences may reconcile diverging interpretations for the role of visual workspace separation as a cue in force fields and cursor rotations (Hegele and Heuer 2010; Howard et al. 2013). In force fields, the visual workspace may be part of the state space representation of implicit learning, thus enabling learners to acquire opposing transformations locally in this state space when visual workspace locations are separate (Howard et al. 2013). In cursor rotations on the other hand, the role of the visual workspace appears confined to being a contextual cue for explicit strategies (Hegele and Heuer 2010), as corroborated by our current findings.

Despite these differences, the recent studies showing that opposing force fields can be learned when different plans are associated with identical trajectories (Hirashima and Nozaki 2012; Sheahan et al. 2016, 2018) indicate that similar principles as the ones we found in this study may apply across kinematic and dynamic transformations. Note that our current results extend these findings by showing that the plan does not need to be tied to a visual target, in line with learning from sensory prediction errors being independent of visual target presence (Lee et al. 2018). To which extent plan-dependent force compensation can be characterized as explicit or implicit remains to be clarified.

### Relation to models of dual adaptation

A number of models have been proposed to formalize the underlying computational principles of dual adaptation to opposing sensorimotor transformations (Ghahramani and Wolpert 1997; Wolpert and Kawato 1998; Thoroughman and Shadmehr 2000; Donchin et al. 2003; Lee and Schweighofer 2009; Lonini et al. 2009; van Dam and Ernst 2015; McDougle et al. 2017). Many of these models share the common feature that a set of independent bases encodes a visuomotor map and that these bases learn from errors depending on how close the movement was to their preferred direction in some relevant cue space. Applying these models to our results, they can in principle explain the local, implicit generalization phenomena we observed via limited local generalization if we use the explicit movement plan as the relevant cue. This implies a serial arrangement where the motor plan feeds into the implicit adaptation model. A previous model comparison study concluded that distinct motor learning processes are arranged in parallel (Lee and Schweighofer 2009). However, they arrived at this conclusion by sequential comparison of different models with the underlying assumptions neither considering locally limited generalization nor specific properties associated with explicit planning. Therefore, their conclusion may not apply when these are taken into account.

### Implications for experiments and real-world tasks

Irrespective of underlying computations or neural processes, our results have important consequences for behavioral experiments and their interpretation with respect to real-world behavior. Thus, our results show that the implicit mechanism that underlies aftereffects can learn opposing visuomotor transformations locally for specific parts of the workspace without forming separate visuomotor maps, provided that contextual cues allow separate aiming strategies to be associated with opposing transformations. This implies that practice with the same set of targets does not guarantee that resulting aftereffects reflect the formation of separate implicit visuomotor maps, nor do similar motor solutions (Hirashima and Nozaki 2012). Instead, posttests may be probing local dents within a single visuomotor map if participants’ aims resemble those during practice. Alternatively, when participants are instructed to aim towards the visual target, aftereffects may be absent because the generalization function is probed at the point of maximal interference, as in our experiments 1 and 2. This phenomenon is also likely to explain absence of aftereffects in a previous study of ours (Hegele and Heuer 2010). Overall, researchers need to take into account the possibility that flexible movement plans form a complex generalization landscape, particularly when learners are practicing multiple sensorimotor transformations.

With respect to the introductory example, we would expect the use of different tools to be realized by distinct motor memories comprising different visuomotor mappings, given that successful tool use does not appear to be constrained to a small range of directions, except by biomechanical constraints. Given our current results and that most studies on dual adaptation do not differentiate whether learning observed by them is explicit or implicit and if it occurs locally or in separate visuomotor maps, we do not currently see compelling evidence that the mechanism that underlies implicit aftereffects in visuomotor dual adaptation is indeed relevant for learning to use different tools by a priori context inference. Further research is needed to identify cues that may indeed support separate, implicit visuomotor maps by this mechanism. Other than identifying “pre-existing” cues, an interesting question is whether such cues to separate implicit learning may be learned by associations over a longer time scale, as suggested previously (Howard et al. 2013). Alternatively, participants could learn these skills by explicit strategies that become automatized into implicit tendencies for action selection (Morehead et al. 2015), in line with canonical theories of motor skill learning (Fitts and Posner 1967). The role of the process that produces aftereffects on the other hand may be limited to calibrating the system to changes that are more biologically common, such as muscular fatigue. Such a division of responsibilities would be reminiscent of the classical distinction between learning of intrinsic (body) vs. extrinsic (tool) transformations (Heuer 1983). Our results therefore highlight the importance of distinguishing between different concepts of dual adaptation, i.e. local shaping of a single versus the formation of separate visuomotor maps.

## Acknowledgements

We thank Lisa M Langsdorf, Simon Koch, Samuel Poggemann, Vanessa Walter and Simon Rosental for data collection. We further thank Eugene Poh and Sam McDougle for helpful discussions on data analysis.

